# Vaccine-breakthrough infection by the SARS-CoV-2 Omicron variant elicits broadly cross-reactive immune responses

**DOI:** 10.1101/2021.12.27.474218

**Authors:** Runhong Zhou, Kelvin Kai-Wang To, Qiaoli Peng, Jacky Man-Chun Chan, Haode Huang, Dawei Yang, Bosco Hoi-Shiu Lam, Vivien Wai-Man Chuang, Jian-Piao Cai, Na Liu, Ka-Kit Au, Owen Tak-Yin Tsang, Kwok-Yung Yuen, Zhiwei Chen

## Abstract

Highly transmissible SARS-CoV-2 Omicron variant has posted a new crisis for COVID-19 pandemic control. Within a month, Omicron is dominating over Delta variant in several countries probably due to immune evasion. It remains unclear whether vaccine-induced memory responses can be recalled by Omicron infection. Here, we investigated host immune responses in the first vaccine-breakthrough case of Omicron infection in Hong Kong. We found that the breakthrough infection rapidly recruited potent cross-reactive broad neutralizing antibodies (bNAbs) against current VOCs, including Alpha, Beta, Gamma, Delta and Omicron, from unmeasurable IC_50_ values to mean 1:2929 at around 9-12 days, which were higher than the mean peak IC_50_ values of BioNTech-vaccinees. Cross-reactive spike- and nucleocapsid-specific CD4 and CD8 T cell responses were detected. Similar results were also obtained in the second vaccine-breakthrough case of Omicron infection. Our preliminary findings may have timely implications to booster vaccine optimization and preventive strategies of pandemic control.

## Main text

Highly transmissible SARS-CoV-2 and its variants have caused more than 279 million infections with about 5.4 million deaths globally by December 26, 2021 (https://coronavirus.jhu.edu/map.html). To fight the ongoing pandemic, 8.9 billion doses of several types of COVID-19 vaccine have already been extensively administered in many countries, which has reduced the rates of hospitalization and death significantly (Baden et al., 2021; Polack et al., 2020; Tanriover et al., 2021; Voysey et al., 2021; Xia et al., 2021). Since these vaccines cannot confer complete prevention of upper airway transmission of SARS-CoV-2, the increasing numbers of vaccine-breakthrough infections and re-infections have been documented (Abu-Raddad et al., 2021; Birhane et al., 2021; To et al., 2021). This situation is becoming worse because of the rapid spread of SARS-CoV-2 variant of concerns (VOCs) and waning of vaccine-induced immune responses (Peng et al., 2021; Wang et al., 2021a; Wang et al., 2021b). After World Health Organization (WHO) designated the Omicron variant of concern (VOC) on the 26^th^ of November 2021, the extremely rapid spread of this variant has led to another crisis of pandemic control. Within a month, Omicron is replacing the Delta VOC to become the dominant SARS-CoV-2 variant in many places in the South Africa, European countries and in the United States (Shu and McCauley, 2017). Two studies reported that the increased risk of re-infection was associated with emergence of Omicron in South Africa and Denmark (Espenhain et al., 2021; Pulliam et al., 2021). Both vaccine-induced neutralizing antibody (NAb) and current NAb combination therapy for passive immunization have significantly reduced activities (Lu et al., 2021; Wang et al., 2021a). Till now, it remains unclear whether vaccine-induced memory responses can be recalled by the Omicron viral infection. We, therefore, investigated the host immune responses in two vaccine-breakthrough cases of Omicron infection in Hong Kong. Our preliminary finding of Omicron-recalled broadly cross-reactive immune responses in these cases may have timely important implications to booster vaccine optimization and implementing adequate preventive interventions to control the pandemic.

On mid-November 2021, the first Chinese vaccine-breakthrough case of Omicron patient (OP1) was diagnosed in a quarantine hotel in Hong Kong (Wong et al., 2021). OP1 arrived in Hong Kong from Canada and was tested negative by reverse transcription PCR (RT-PCR) for SARS-CoV-2 within 72 hours before arrival. Seven days after arrival, OP1 developed mild symptoms and showed a positive result for SARS-CoV-2 (Ct value 19) on day 8 after arrival and was hospitalized on the same day. To validate our findings, we subsequently received blood samples from an imported mild case of Omicron patient 2 (OP2), who was due to a separate transmission and was diagnosed about 9 days after the OP1. Based on the vaccination records, OP1 and OP2 were confirmed with Omicron infection at 178 and 53 days after the second dose of BNT162b2 and mRNA-1273, respectively (**Figure 1A**). During hospitalization, both cases presented with mild clinical symptoms not requiring oxygen supplementation or ICU treatment. With patients’ informed consent, we obtained three sequential sera and one peripheral blood mononuclear cells (PBMCs) samples from each patient to determine whether vaccine-induced memory responses can be recalled by the Omicron viral infection.

**Figure 1.**
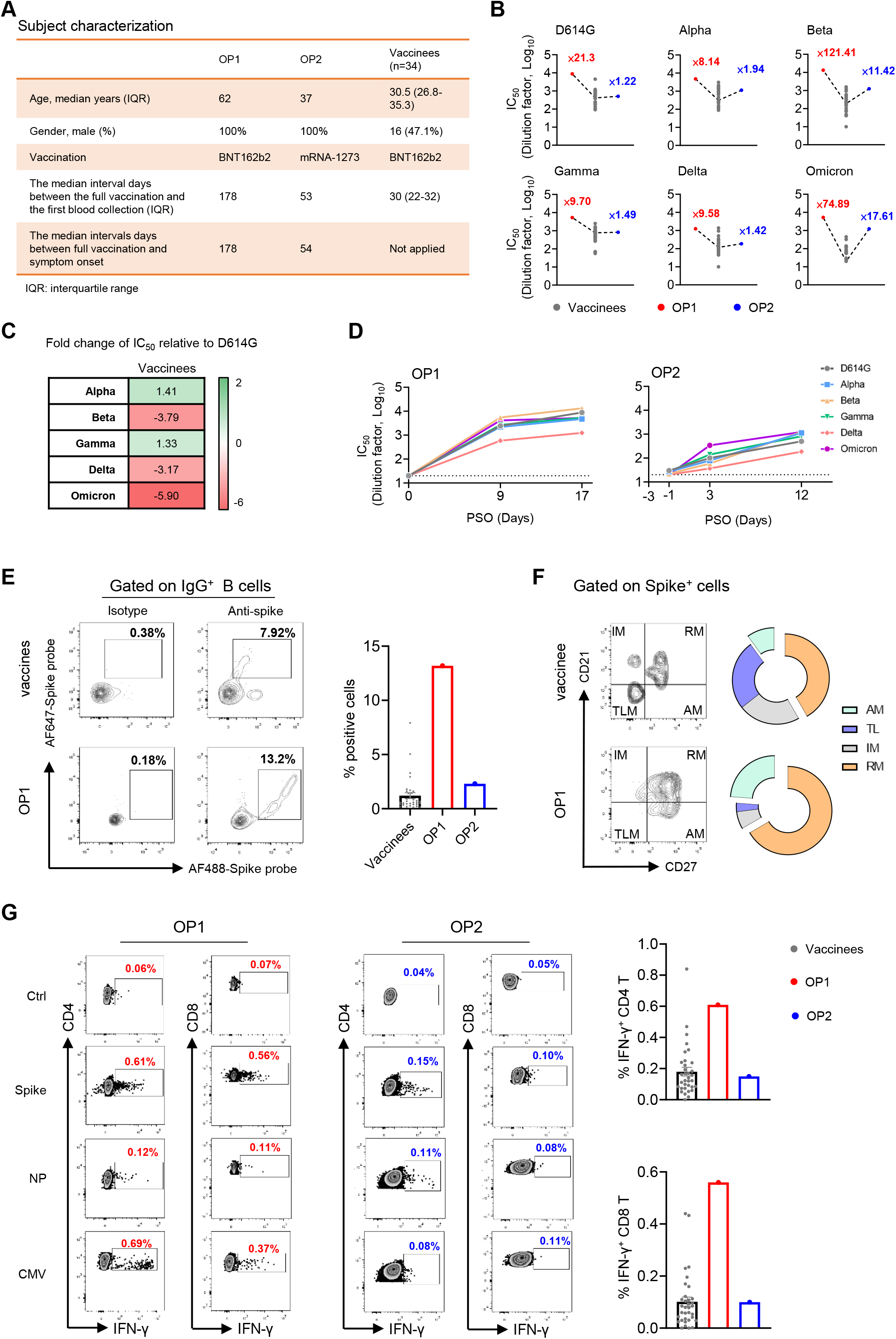
Cross-reactive immune responses elicited by vaccine-breakthrough infection of the SARS-CoV-2 Omicron variant. (**A**) Characterization of 2 Omicron patients and 34 BNT162b2-vaccinees. (**B**) Neutralizing antibody titers among the BNT162b2-vaccinees (grey) (n=34) and two Omicron patients (OP1: red and OP2: blue) at the peak response time. Neutralizing antibody titers represent serum dilution required to achieve 50% virus neutralization (IC_50_). The numbers indicate the fold of enhancement of IC_50_ values relative to mean titer measured among BNT162b2-vaccinees. (**C**) Fold-change of mean IC_50_ values relative to the SARS-CoV-2 D614G strain among the BNT162b2-vaccinees. (**D**) Longitudinal neutralizing antibody titers (IC_50_) of OP1 and OP2 against the full panel of VOCs. Each symbol with color-coding represents an individual VOC. (**E**) The gating strategy for SARS-CoV-2 Spike-specific B cells by flow cytometry. AF488 and AF647 double positive cells were defined as Spike-specific cells. Representative plots (left) and quantified results (right) are shown. (**F**) Phenotypes of Spike-specific B cells were defined by using CD21 and CD27 markers (left). Pie chart showed the proportion of activated (AM), tissue-like memory (TLM), intermediate memory (IM) and resting-memory (RM) B cells. (**G**) PBMCs were subjected to the ICS assay against Spike or NP or CMV peptide pools. IFN-γ^+^ cells were gated on CD4 and CD8 T cells, respectively (left). Quantified results (right) depict the percentage of IFN-γ^+^ cells.

We first measured the neutralizing antibody titer (IC_50_) in their sera samples against the current panel of SARS-CoV-2 VOC pseudoviruses including Alpha (B.1.1.7), Beta (B. 1.351), Gamma, Delta (B. 1.617.2) and Omicron (B. 1.1.529) as compared with D614G (WT). We compared IC_50_ values with 34 local vaccinees, whose blood was collected around mean 30 days after the second BNT162b2-vaccination (Pfizer–BioNTech) (**Figure 1A**) (Peng et al., 2021). Consistent with recent preprint publications by others, we found that the Omicron variant showed the greatest resistance to BNT162b2-vaccine-induced neutralization with an average 5.9-fold deficit relative to D614G (**Figure 1C**). Strikingly, however, the breakthrough infection was able to elicit cross-reactive broad neutralizing antibodies (bNAbs) from the unmeasurable levels (<1:20) to mean IC_50_ values of 1:2929 (range 588.5-5508) and from mean IC_50_ 1:24.3 to 1:854.5 at 9 days in OP1 and 12 days in OP2 post symptoms onset (PSO), respectively (**Figure 1D**). Moreover, the amounts of NAbs were consistently higher than the mean IC_50_ values of BNT162b2-vaccinees across all VOCs tested. In particular, there were 121.41- and 74.89-fold higher IC_50_ values against Beta and Omicron in OP1 than those in BNT162b2-vaccinees (**Figure 1B**). Besides NAbs against the current panel of VOCs, OP1 also displayed enhanced IC_50_ values of NAbs against 15/16 SARS-CoV-2 variants with individual mutations or deletions including the E484K mutation, which conferred significant resistance to vaccine-induced NAbs (**Figure S1**). These results demonstrated that, although the Omicron VOC evaded BNT162b2-vaccine-induced NAbs, the breakthrough infection elicited cross-reactive bNAbs generally against all current VOCs in both OP1 and OP2.

To understand cellular immune responses, we conducted flow cytometry analysis on PBMCs of OP1 and OP2 collected on day 11 and 12 PSO, respectively. Multi-color flow cytometry data showed no sign of severe immune suppression in both OP1 and OP2 who had similar frequencies of T lymphocyte without lymphocytopenia, stable conventional dendritic cell (cDC): plasmacytoid dendritic cell (pDC) ratio and normal Myeloid-derived suppressor cells (MDSCs) to mild and healthy subjects as we described previously (**Figure S2**) (Zhou et al., 2020). For antigen-specific B cell activation, we measured the frequency of Spike-specific IgG^+^ B cells in OP1 and OP2. The levels of 13.2% in OP1 and 2.31% in OP2 were relatively higher than mean 1.12% (range 0.004-7.92%) found among BNT162b2-vaccinees around their peak responses (**Figure 1E**). Unlike naturally infected COVID-19 patients, who display predominantly tissue-like memory (TLM) B cell response (Woodruff et al., 2020), Spike-specific IgG^+^ B cells from OP1 and OP2 exhibited the dominate phenotype of resting memory (RM) (**Figure 1F**), which was also found in our BNT162b2-vaccinees.

Besides Spike-specific IgG^+^ B cell responses, we measured cross-reactive T cell responses to the Spike and nucleocapsid (NP) peptide pools derived from wildtype SARS-CoV-2 in OP1 and OP2 as compared with BNT162b2-vaccinees by intracellular cytokine staining. The cytomegalovirus (CMV) pp65 peptide pool was used as a positive control. We found that Spike- and NP-specific CD4 IFN-γ responses were 0.61% and 0.12% in OP1 and 0.15% and 0.10% in OP2, respectively (**Figure 1G**). Moreover, Spike- and NP-specific CD8 IFN-γ responses were 0.56% and 0.11% in OP1 and 0.10% and 0.08% in OP2, respectively (**Figure 1G**). These results indicated that cross-reactive CD4 and CD8 T cell responses to wild type SARS-CoV-2 were primarily against the Spike as compared with NP. Moreover, the Spike-specific T cell responses were relatively higher in OP1 or comparable in OP2 as compared with mean values in BNT162b2-vaccinees (CD4 T: mean 0.19% and CD8 T: mean 0.10%). Since much weaker or unmeasurable T cell responses were found in severe COVID-19 patients around the same period PSO (Rydyznski Moderbacher et al., 2020; Zhou et al., 2020), T cell responses in OP1 and OP2 probably also contributed to disease progression control.

In this brief report, we provide timely communication on immune responses in two cases of vaccine-breakthrough infections by the SARS-CoV-2 Omicron variant in Hong Kong. Although antibody evasion has been clearly documented against Omicron due to 32 amino acid changes in viral spike protein (Cameroni et al., 2021; Cele et al., 2021; Planas et al., 2021a; Wang et al., 2021a), we report here that Omicron vaccine-breakthrough infections could elicit cross-reactive bNAb responses against all current SARS-CoV-2 VOCs. Since the amounts of bNAb responses were higher than the mean IC_50_ values of in BNT162b2-vaccinees at their peak response period, we believe that the Omicron infection rapidly recruited the vaccine-induced memory immune responses during the acute phase of infection, which probably contributed to protection and was in line with the mild clinical presentation in both patients. Encouragingly, besides rapid bNAb responses, both spike- and NP-specific CD4 and CD8 T cells cross-reactive to wild type peptide pools were measurable on day 11-12, which probably also contributed to disease progression control (Lipsitch et al., 2020; Zhou et al., 2020).

Both OP1 and OP2 showed high amounts of bNAbs against the Omicron variant and other VOCs. According to the GASAID database, during the period from October 4, 2021 to December 26, 2021, the relative variant genome frequency of the current circulating Delta variant has declined from 89% to 19.6% while the Omicron variant has increased from 0% to 67.4% in African countries. Besides insufficient vaccination coverage and preventive masking, high viral infectivity and antibody escape are likely the key reasons for the rapidity of Omicron spread. Based on *in vitro* experiments, the Omicron variant showed a 10-fold increase in infectivity than the Beta or Delta variants (Lu et al., 2021). Consistent with previous findings that the Beta variant compromised vaccine-induced neutralizing activity (Planas et al., 2021b; Wang et al., 2021a), similar findings have already been made for the Omicron variant with even worse antibody evasion (Cameroni et al., 2021; Cele et al., 2021; Planas et al., 2021a; Wang et al., 2021a). We also made similar findings that a significant drop of neutralizing activity against Omicron variant was observed among the convalescent patients and vaccine recipients (Lu et al., 2021; Wang et al., 2022). Since the Omicron variant caused a higher rate of vaccine-breakthrough infection and reinfection than the Delta variant (Espenhain et al., 2021), it is worrisome if such infections would lead to more severe sickness or death due to immune escape. In this study, we demonstrated that the Omicron breakthrough infection rapidly recruited vaccine-induced memory bNAbs and T cell immune responses, which very likely contributed to protection to both OP1 and OP2. Our finding is consistent with and provides a probable immune mechanism underlying a recent report that most Omicron patients had no signs of severe COVID-19 as compared with the Delta variant (Espenhain et al., 2021). Future studies, however, remain necessary to evaluate Omicron-specific T cell immunity for protection although there were no significant reductions in CD4 and CD8 T cell responses to the spike peptides-derived from Alpha and Delta spike variants (Jordan et al., 2021). Our findings, therefore, re-emphasize the importance of complete vaccination coverage among human populations especially in developing countries. Since similarly high amounts of bNAbs against both Omicron and other VOCs were detected in both OP1 and OP2, the rapid development of Omicron-based vaccine is a reasonable strategy for the booster vaccine optimization.

The major limitation of this study is the small number of vaccine-breakthrough infections by the SARS-CoV-2 Omicron variant found in Hong Kong. Some mutations in Omicron spike are shared with preexisting VOCs, such as D614G in all VOCs, K417N, E484K and N501Y in Beta variant, and T478K in Delta variant. The E484K mutation in Beta variant has been reported for evasion of many NAbs under clinical development (Wang et al., 2021a). These mutations in combination with additional mutations have led to the striking antibody evasion manifested by the Omicron variant (Wang et al., 2021a). Nevertheless, our preliminary finding, that OP1 and OP2 could generate bNAbs against all VOCs after infection, suggested that the Omicron-targeted vaccine might boost a broad protection among existing vaccinees against SARS-CoV-2 VOC infection. Since current vaccines showed weak effect on Omicron, our findings also implicate that the development of Omicron-targeted vaccines is urgent and beneficial to fight all current SARS-CoV-2 VOCs, especially when the increased infectivity of Omicron variant has been preliminarily reported *in vitro* (Lu et al., 2021).

## SUPPLEMENTAL INFORMATION

2 supplemental figures.

## STAR METHODS

### RESOURCE AVAILABILITY

#### Lead Contact

Further information and requests for resources and reagent should be directed to and will be fulfilled by the Lead Contact, Zhiwei Chen.

#### Materials Availability

This study did not generate new unique reagents.

#### Data and Code Availability

The study did not generate any unique datasets or codes.

### EXPERIMENTAL MODELS AND SUBJECT DETAILS

#### Human subjects

This study was approved by the Institutional Review Board of the University of Hong Kong/Hospital Authority Hong Kong West Cluster (Ref No. UW 21-452). Written informed consent was obtained from all study subjects. Peripheral blood mononuclear cells (PBMCs) from healthy donors and patients were isolated from fresh blood samples using Ficoll-Paque density gradient centrifugation in our BSL-3 laboratory at the same day of blood collection. The majority of purified PBMCs were used for immune cell phenotyping whereas plasma samples were subjected to antibody testing.

The rest of the cells were cryopreserved in freezing medium (Synth-a-Freeze Cryopreservation Medium, ThermoFisher Scientific) at 5 × 10^6^ cells/mL at −150°C.

#### Pseudotyped viral neutralization assay

To determine the neutralizing activity of subject’ plasma, plasma was inactivated at 56°C for 30 min prior to a pseudotyped viral entry assay. In brief, different SARS-CoV-2 pseudotyped viruses were generated through co-transfection of 293T cells with 2 plasmids, pSARS-CoV-2 S and pNL4-3Luc_Env_Vpr, carrying the optimized SARS-CoV-2 S gene and a human immunodeficiency virus type 1 backbone, respectively. At 48 h post-transfection, viral supernatant was collected and frozen at −150°C. Serially diluted plasma samples (from 1:20 to 1:14580) were incubated with 200 TCID50 of pseudovirus at 37°C for 1 h. The plasma-virus mixtures were then added into pre-seeded HEK293T-hACE2 cells. After 48 h, infected cells were lysed, and luciferase activity was measured using Luciferase Assay System kits (Promega) in a Victor3-1420 Multilabel Counter (PerkinElmer). The 50% inhibitory concentrations (IC_50_) of each plasma specimen were calculated to reflect anti-SARS-CoV-2 potency.

#### Flow cytometry analysis

For immune cell profile analysis, PBMCs were incubated for 10 min with Fc Block (BD Biosciences) in staining buffer (PBS containing 2% FBS) followed by staining with the indicated antibodies for 30 min at 4°C. For T cell responses, PBMCs were stimulated with 2 μg/mL COVID-19 Spike or NP peptide pool (15-mer overlapping by 11) or CMV pp65 peptide pool in the presence of 0.5 μg/mL anti-CD28 and anti-CD49d mAbs (BD Bioscience). Cells were incubated at 37°C overnight and BFA was added at 2 h post incubation, as previously described (Li et al., 2008a). After overnight incubation, cells were washed with staining buffer (PBS containing 2% FBS) and stained with mAbs against surface markers. For intracellular staining, cells were fixed and permeabilized with BD Cytofix/Cytoperm (BD Biosciences) prior to staining with the mAbs against cytokines with Pern/Wash buffer (BD Biosciences). Stained cells were acquired by FACSAriaIII Flow Cytometer (BD Biosciences) inside a BSL-3 laboratory and analyzed with FlowJo software (v10.6) (BD Bioscience).

## ACKNOWLEDGMENTS

This study was supported by the Hong Kong Research Grants Council Collaborative Research Fund (C7156-20G, C1134-20G and C5110-20G) and Health and Medical Research Fund (19181012); Shenzhen Science and Technology Program (JSGG20200225151410198 and JCYJ20210324131610027); the Hong Kong Health@InnoHK, Innovation and Technology Commission; and the China National Program on Key Research Project (2020YFC0860600, 2020YFA0707500 and 2020YFA0707504); and donations from the Friends of Hope Education Fund. Z.C.’s team was also partly supported by the Hong Kong Theme-Based Research Scheme (T11-706/18-N). We sincerely thank Drs. David D. Ho and Pengfei Wang for the expression plasmids encoding for D614G, Alpha and Beta variants, Dr. Linqi Zhang for the Delta variant plasmid.

## AUTHOR CONTRIBUTIONS

Z.C. and R.Z conceived and designed the study. R.Z. and Z.C. designed experiments, analyzed data, and wrote the manuscript. R.Z., H.H., and N.L. performed the flow cytometry analysis. K.K.-W.T., J.M.-C.C., B.H-.S.L., V.W.-M.C. and O.T.-Y.T. collected clinical samples and data. Q.P., D.Y. and K.-K.A conducted the pseudoviral neutralization assay. J.-P.C. provided technique support. K.-Y.Y. provided critical comments.

## References

Abu-Raddad, L.J., Chemaitelly, H., and Bertollini, R. (2021). Severity of SARS-CoV-2 Reinfections as Compared with Primary Infections. New Engl J Med.

Baden, L.R., El Sahly, H.M., Essink, B., Kotloff, K., Frey, S., Novak, R., Diemert, D., Spector, S.A., Rouphael, N., Creech, C.B., et al. (2021). Efficacy and Safety of the mRNA-1273 SARS-CoV-2 Vaccine. N Engl J Med 384, 403–416.

Birhane, M., Bressler, S., Chang, G., Clark, T., Dorough, L., Fischer, M., Watkins, L.F., Goldstein, J.M., Kugeler, K., Langley, G., et al. (2021). COVID-19 Vaccine Breakthrough Infections Reported to CDC — United States, January 1–April 30, 2021. MMWR Morb Mortal Wkly Rep 70, 792–793.

Cameroni, E., Bowen, J., and Corti, D. (2021). Broadly neutralizing antibodies overcome SARS-CoV-2 Omicron antigenic shift. Nature https://doi.org/10.1038/d41586-021-03825-4.

Cele, S., Jackson, L., and Sigal, A. (2021). Omicron extensively but incompletely escapes Pfizer BNT162b2 neutralization. Nature https://doi.org/10.1038/d41586-021-03824-5.

Espenhain, L., Funk, T., Overvad, M., Edslev, S.M., Fonager, J., Ingham, A.C., Rasmussen, M., Madsen, S.L., Espersen, C.H., Sieber, R.N., et al. (2021). Epidemiological characterisation of the first 785 SARS-CoV-2 Omicron variant cases in Denmark, December 2021. Euro Surveill 26.

Jordan, S.C., Shin, B.H., Gadsden, T.M., Chu, M., Petrosyan, A., Le, C.N., Zabner, R., Oft, J., Pedraza, I., Cheng, S., et al. (2021). T cell immune responses to SARS-CoV-2 and variants of concern (Alpha and Delta) in infected and vaccinated individuals. Cell Mol Immunol 18, 2554–2556.

Lipsitch, M., Grad, Y.H., Sette, A., and Crotty, S. (2020). Cross-reactive memory T cells and herd immunity to SARS-CoV-2. Nat Rev Immunol 20, 709–713.

Lu, L., Mok, B.W., Chen, L.L., Chan, J.M., Tsang, O.T., Lam, B.H., Chuang, V.W., Chu, A.W., Chan, W.M., Ip, J.D., et al. (2021). Neutralization of SARS-CoV-2 Omicron variant by sera from BNT162b2 or Coronavac vaccine recipients. Clin Infect Dis.

Peng, Q., Zhou, R., Wang, Y., Zhao, M., Liu, N., Li, S., Huang, H., Yang, D., Au, K.-K., Wang, H., et al. (2021). Waning immune responses against SARS-CoV-2 among vaccinees in Hong Kong.

Planas, D., Saunders, N., and Schwartz, O. (2021a). Considerable escape of SARS-CoV-2 Omicron to antibody neutralization. Nature https://doi.org/10.1038/d41586-021-03827-2.

Planas, D., Veyer, D., Baidaliuk, A., Staropoli, I., Guivel-Benhassine, F., Rajah, M.M., Planchais, C., Porrot, F., Robillard, N., Puech, J., et al. (2021b). Reduced sensitivity of SARS-CoV-2 variant Delta to antibody neutralization. Nature 596, 276–280.

Polack, F.P., Thomas, S.J., Kitchin, N., Absalon, J., Gurtman, A., Lockhart, S., Perez, J.L., Perez Marc, G., Moreira, E.D., Zerbini, C., et al. (2020). Safety and Efficacy of the BNT162b2 mRNA Covid-19 Vaccine. N Engl J Med 383, 2603–2615.

Pulliam, J.R.C., van Schalkwyk, C., Govender, N., von Gottberg, A., Cohen, C., Groome, M.J., Dushoff, J., Mlisana, K., and Moultrie, H. (2021). Increased risk of SARS-CoV-2 reinfection associated with emergence of the Omicron variant in South Africa. medRxiv, 2021.2011.2011.21266068.

Rydyznski Moderbacher, C., Ramirez, S.I., Dan, J.M., Grifoni, A., Hastie, K.M., Weiskopf, D., Belanger, S., Abbott, R.K., Kim, C., Choi, J., et al. (2020). Antigen-Specific Adaptive Immunity to SARS-CoV-2 in Acute COVID-19 and Associations with Age and Disease Severity. Cell 183, 996–1012 e1019.

Shu, Y., and McCauley, J. (2017). GISAID: Global initiative on sharing all influenza data - from vision to reality. Euro Surveill 22.

Tanriover, M.D., Doganay, H.L., Akova, M., Guner, H.R., Azap, A., Akhan, S., Kose, S., Erdinc, F.S., Akalin, E.H., Tabak, O.F., et al. (2021). Efficacy and safety of an inactivated whole-virion SARS-CoV-2 vaccine (CoronaVac): interim results of a double-blind, randomised, placebo-controlled, phase 3 trial in Turkey. Lancet 398, 213–222.

To, K.K., Hung, I.F., Ip, J.D., Chu, A.W., Chan, W.M., Tam, A.R., Fong, C.H., Yuan, S., Tsoi, H.W., Ng, A.C., et al. (2021). Coronavirus Disease 2019 (COVID-19) Re-infection by a Phylogenetically Distinct Severe Acute Respiratory Syndrome Coronavirus 2 Strain Confirmed by Whole Genome Sequencing. Clin Infect Dis 73, e2946–e2951.

Voysey, M., Clemens, S.A.C., Madhi, S.A., Weckx, L.Y., Folegatti, P.M., Aley, P.K., Angus, B., Baillie, V.L., Barnabas, S.L., Bhorat, Q.E., et al. (2021). Safety and efficacy of the ChAdOx1 nCoV-19 vaccine (AZD1222) against SARS-CoV-2: an interim analysis of four randomised controlled trials in Brazil, South Africa, and the UK. Lancet 397, 99–111.

Wang, P., Nair, M.S., Liu, L., Iketani, S., Luo, Y., Guo, Y., Wang, M., Yu, J., Zhang, B., Kwong, P.D., et al. (2021a). Antibody resistance of SARS-CoV-2 variants B.1.351 and B.1.1.7. Nature 593, 130–135.

Wang, Y., Chen, R., Hu, F., Lan, Y., Yang, Z., Zhan, C., Shi, J., Deng, X., Jiang, M., Zhong, S., et al. (2021b). Transmission, viral kinetics and clinical characteristics of the emergent SARS-CoV-2 Delta VOC in Guangzhou, China. EClinicalMedicine 40, 101129.

Wang, Y., Zhang, L., Li, Q., Liang, Z., Li, T., Liu, S., Cui, Q., Nie, J., Wu, Q., Qu, X., and Huang, W. (2022). The significant immune escape of pseudotyped SARS-CoV-2 variant Omicron. Emerg Microbes Infect 11, 1–5.

Wong, S.C., Au, A.K., and Cheng, V.C. (2021). Transmission of Omicron (B.1.1.529) - SARS-CoV-2 Variant of Concern in a designated quarantine hotel for travelers: a challenge of elimination strategy of COVID-19. The Lancet Regional Health - Western Pacific https://doi.org/10.1016/j.lanwpc.2021.100360.

Woodruff, M.C., Ramonell, R.P., Nguyen, D.C., Cashman, K.S., Saini, A.S., Haddad, N.S., Ley, A.M., Kyu, S., Howell, J.C., Ozturk, T., et al. (2020). Extrafollicular B cell responses correlate with neutralizing antibodies and morbidity in COVID-19. Nat Immunol 21, 1506–1516.

Xia, S., Zhang, Y., Wang, Y., Wang, H., Yang, Y., Gao, G.F., Tan, W., Wu, G., Xu, M., Lou, Z., et al. (2021). Safety and immunogenicity of an inactivated SARS-CoV-2 vaccine, BBIBP-CorV: a randomised, double-blind, placebo-controlled, phase 1/2 trial. Lancet Infect Dis 21, 39–51.

Zhou, R., To, K.K., Wong, Y.C., Liu, L., Zhou, B., Li, X., Huang, H., Mo, Y., Luk, T.Y., Lau, T.T., et al. (2020). Acute SARS-CoV-2 Infection Impairs Dendritic Cell and T Cell Responses. Immunity 53, 864–877 e865.

