## Supplemental figures for "Vaccine-breakthrough infection by the SARS-CoV-2 Omicron variant elicits broadly cross-reactive immune responses": Supplementary Figures.pdf

| Individual mutations/deletion | $IC_{50}$ values | Fold change |
|---|---|---|
| D614G | 1521 | 1 |
| Del 69-70 | 5463 | 3.6 |
| Del 144 | 4989 | 3.3 |
| N501Y | 5606 | 3.7 |
| A570D | 4661 | 3.1 |
| P681H | 8038 | 5.3 |
| T716I | 2106 | 1.4 |
| S982A | 13506 | 8.9 |
| D1118H | 8007 | 5.3 |
| L18F | 6084 | 4.0 |
| D80A | 2895 | 1.9 |
| D215G | 1552 | 1.0 |
| Del242-244 | 3409 | 2.2 |
| R246I | 7324 | 4.8 |
| K417N | 2864 | 1.9 |
| E484K | 4198 | 2.8 |
| A701V | 5858 | 3.9 |

**Figure S1**. Plasma sample was collected at 13 days after symptom onset of OP1 and was subjected to the neutralizing activity test against the pseudoviruses containing individual mutation or deletion. Neutralizing antibody titers represent serial dilution that achieved 50% virus neutralization ($IC_{50}$). The fold change of $IC_{50}$ values was relative to the D614G variant.

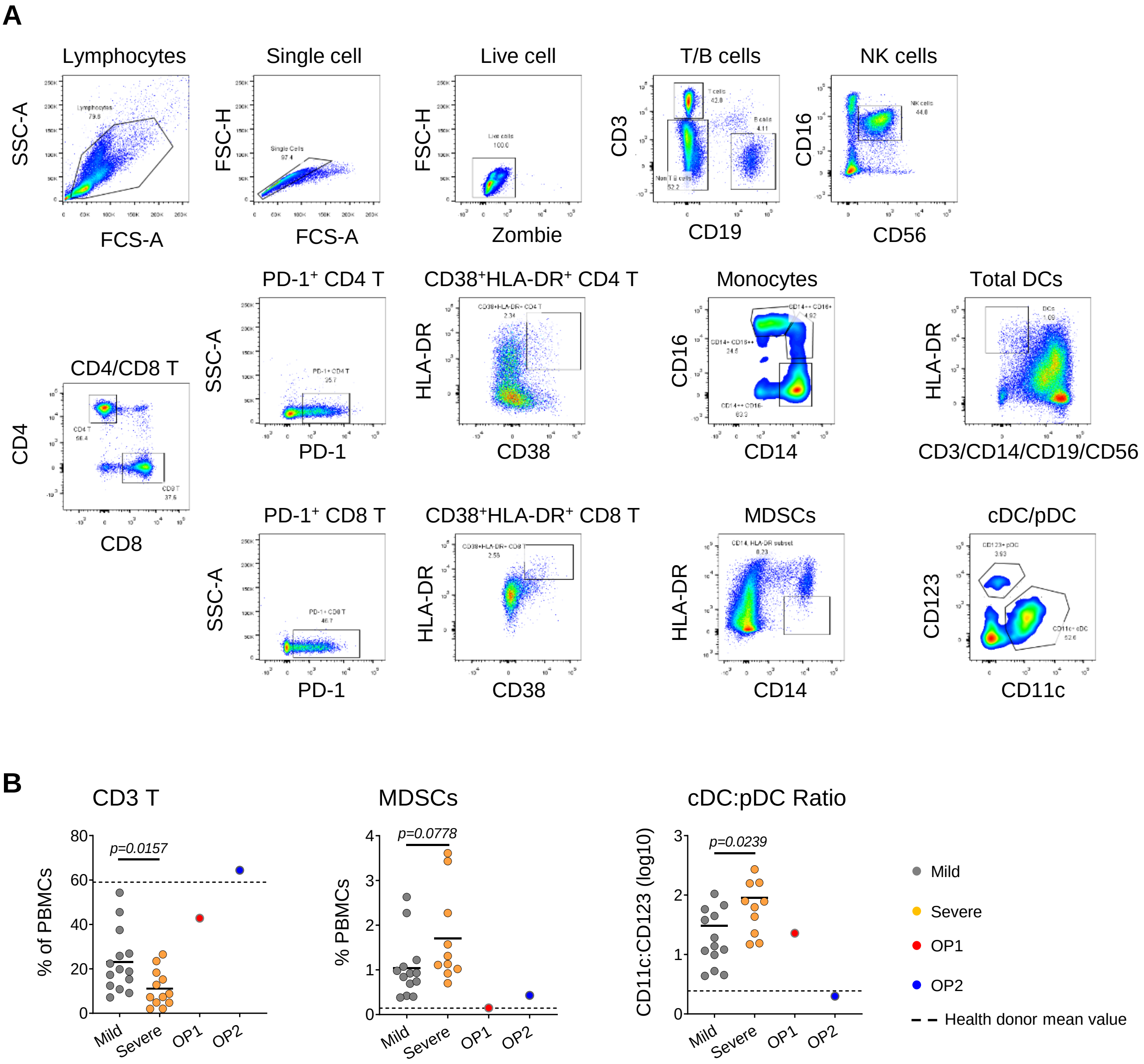


**Figure S2**. (**A**) Gating strategy of immune cell profile. (**B**) Cumulative data show various cell frequencies. Each symbol represents a study subject, and the solid line indicates the mean of each group. Mild: mild patients. Severe: Severe patients. The dash line indicates the mean value of health donors. Statistics were generated by using 2-tailed Student's t test.
